# Illumina TruSeq synthetic long-reads empower *de novo* assembly and resolve complex, highly repetitive transposable elements

**DOI:** 10.1101/001834

**Authors:** Rajiv C. McCoy, Ryan W. Taylor, Timothy A. Blauwkamp, Joanna L. Kelley, Michael Kertesz, Dmitry Pushkarev, Dmitri A. Petrov, Anna-Sophie Fiston-Lavier

## Abstract

High-throughput DNA sequencing technologies have revolutionized genomic analysis, including the *de novo* assembly of whole genomes. Nevertheless, assembly of complex genomes remains challenging, in part due to the presence of dispersed repeats which introduce ambiguity during genome reconstruction. Transposable elements (TEs) can be particularly problematic, especially for TE families exhibiting high sequence identity, high copy number, or present in complex genomic arrangements. While TEs strongly affect genome function and evolution, most current *de novo* assembly approaches cannot resolve long, identical, and abundant families of TEs. Here, we applied a novel Illumina technology called TruSeq synthetic long-reads, which are generated through highly parallel library preparation and local assembly of short read data and achieve lengths of 1.5-18.5 Kbp with an extremely low error rate (*∼*0.03% per base). To test the utility of this technology, we sequenced and assembled the genome of the model organism *Drosophila melanogaster* (reference genome strain *y;cn,bw,sp*) achieving an N50 contig size of 69.7 Kbp and covering 96.9% of the euchromatic chromosome arms of the current reference genome. TruSeq synthetic long-read technology enables placement of individual TE copies in their proper genomic locations as well as accurate reconstruction of TE sequences. We entirely recovered and accurately placed 4,229 (77.8%) of the 5,434 of annotated transposable elements with perfect identity to the current reference genome. As TEs are ubiquitous features of genomes of many species, TruSeq synthetic long-reads, and likely other methods that generate long reads, offer a powerful approach to improve *de novo* assemblies of whole genomes.

## Introduction

Tremendous advances in DNA sequencing technology, computing power, and assembly approaches, have enabled the assembly of genomes of thousands of species from the sequences of DNA fragments, but several challenges still remain. All assembly approaches are based on the assumption that similar sequence reads originate from the same genomic region, thereby allowing the reads to be overlapped and merged to reconstruct the underlying genome sequence [1]. Deviations from this assumption, including those arising due to polymorphism and repeats, complicate assembly and may induce assembly failure. When possible, performing multiple rounds of inbreeding, using input DNA from a single individual, or even sequencing mutant haploid embryos [2] can limit heterozygosity and improve assembly results.

By spanning regions of high diversity and regions of high identity, the use of longer input sequences can also help overcome problems posed by both polymorphism and repeats. The recent application of Pacific Biosciences (PacBio) long read technology to resolve complex segmental duplications [3] is a case in point. Illumina recently introduced TruSeq™ synthetic long-read technology, which builds upon underlying short read data to generate accurate synthetic reads up to 18.5 Kbp in length. The technology was already used for the *de novo* assembly of the genome of the colonial tunicate, *Botryllus schlosseri* [4]. However, because no high quality reference genome was previously available for that species, advantages, limitations, and general utility of the technology for genome assembly were difficult to assess. By performing assembly of the *Drosophila melanogaster* genome, our study uses comparison to a high quality reference to evaluate the application of synthetic long-read technology for *de novo* assembly. While future work will be required to investigate the use of the technology for resolving polymorphism in outbred species, our work specifically focuses on the accuracy of assembly of repetitive DNA sequences.

In some species, repetitive DNA accounts for a large proportion of the total genome size, for example comprising more than half of the human genome [5, 6] and 80% of some plant genomes [7]. Here, we focus on one class of dynamic repeats, called transposable elements (TEs), which are a common feature of almost all eukaryotic genomes sequenced to date. Some families of TEs are represented in hundreds or even thousands of nearly-identical copies, and some copies span up to tens of kilobases. Consequently, TEs dramatically affect genome size and structure, as well as genome function; transposition has the potential to induce complex genomic rearrangements that detrimentally affect the host, but can also provide the raw material for adaptive evolution [8–10], for example, by creating new transcription factor binding sites [11] or otherwise affecting expression of nearby genes [12].

Despite their biological importance, knowledge of TE dynamics is hindered by technical limitations resulting in the absence of certain TE families from genome assemblies. Many software packages for whole genome assembly use coverage-based heuristics, distinguishing putative unique regions from putative repetitive regions based on deviation from average coverage (e.g., Celera [13], Velvet [14]). While TE families with sufficient divergence among copies may be properly assembled, recently diverged families are often present in sets of disjointed reads or small contigs that cannot be placed with respect to the rest of the assembly. For example, the *Drosophila* 12 Genomes Consortium [15] did not even attempt to evaluate accuracy or completeness of TE assembly. Instead, they used four separate programs to estimate abundance of TEs and other repeats within each assembled genome, but the resulting upper and lower bounds commonly differed by more than three fold. The recent improvement to the draft assembly of *Drosophila simulans* reported that the majority of TE sequences (identified by homology to *D. melanogaster* TEs) were contained in fragmented contigs less than 500 bp in length [16].

TEs, as with other classes of repeats, may also induce mis-assembly. For example, TEs that lie in tandem may be erroneously collapsed, and unique interspersed sequences may be left out or appear as isolated contigs. Several studies have assessed the impact of repeat elements on *de novo* genome assembly. For example, Alkan et al. [17] showed that the human assemblies are on average 16.2% shorter than expected, mainly due to failure to assemble repeats, especially TEs and segmental duplications. A similar observation was made for the chicken genome, despite the fact that repeat density in this genome is lower than humans [18]. In addition to coverage, current approaches to deal with repeats such as TEs generally rely on paired-end data [17, 19, 20]. Paired-end reads can help resolve the orientation and distance between assembled flanking sequences, and repeat-containing reads can sometimes be placed based on uniquely anchored mates. However, if read pairs do not completely span an identical repeat so that at least one read is anchored in unique sequence, alternative possibilities for contig extension cannot be ruled out. Long inserts, commonly referred to as mate-pair libraries, are therefore useful to bridge across long TEs to link and orient contigs, but produce stretches of unknown sequence.

A superior way to resolve TEs is to generate reads that exceed TE length, obviating assembly and allowing TEs to be unambiguously placed based on unique flanking sequence. Pacific Biosciences (PacBio) represents the only high throughput long read (up to *∼*15 Kbp) technology available to date, though Oxford Nanopore [21] platforms may soon be available. While single-pass PacBio sequencing has a high error rate of 15-18%, multiple-pass circular consensus sequencing [22] and hybrid or self error correction [23] improve read accuracy to greater than 99.9%. Meanwhile, other established sequencing technologies, such as Illumina, 454 (Roche), and Ion Torrent (Life Technologies), offer high throughput and low error rates of 0.1-1%, but much shorter read lengths [24]. Illumina TruSeq synthetic long-reads, which are assembled from underlying Illumina short read data, achieve lengths and error rates comparable to PacBio corrected sequences, but their utility for *de novo* assembly has yet to be demonstrated in cases where a high quality reference genome is available for comparison.

Using a pipeline of standard existing tools, we demonstrate the ability of TruSeq synthetic long-reads to facilitate *de novo* assembly and resolve TE sequences in the genome of the fruit fly *Drosophila melanogaster*, a key model organism in both classical genetics and molecular biology. We further investigate how coverage of synthetic long-reads affects assembly results, an important practical consideration for experimental design. While the *D. melanogaster* genome is moderately large (*∼*180 Mbp) and complex, it has already been assembled to unprecedented accuracy. Through a massive collaborative effort, the initial genome project [25] recovered nearly all of the 120 Mbp euchromatic sequence using a whole-genome shotgun approach that involved painstaking molecular cloning and the generation of a bacterial artificial chromosome physical map. Since that publication, the reference genome has been extensively annotated and improved using several resequencing, gap-filling, and mapping strategies, and currently represents a gold standard for the genomics community [26–28]. By performing the assembly in this model system with a high quality reference genome, our study is the first to systematically document the advantages and limitations posed by this synthetic long-read technology. *D. melanogaster* harbors a large number (*∼*100) of families of active TEs, some of which contain many long and virtually identical copies distributed across the genome, thereby making their assembly a particular challenge. This is distinct from other species, including humans, which have TE copies that are shorter and more diverged from each other, and therefore easier to assemble. Our demonstration of accurate TE assembly in *D. melanogaster* should therefore translate favorably to many other systems.

## Results

### TruSeq synthetic long-reads

#### Library preparation

This study used Illumina TruSeq synthetic long-read technology generated with a novel highly-parallel next-generation library preparation method (Figure S1). The basic protocol was previously presented by Voskoboynik et al. [4] (who referred to it as LR-seq) and was patented by Stanford University and licensed to Moleculo, which was later acquired by Illumina. The protocol (see Methods) involves initial mechanical fragmentation of gDNA into *∼*10 Kbp fragments. These fragments then undergo end-repair and ligation of amplification adapters, before being diluted onto 384-well plates so that each well contains DNA representing approximately 1-2% of the genome (*∼*200 molecules, in the case of *Drosophila melanogaster)*. Polymerase chain reaction (PCR) is used to amplify molecules within wells, followed by highly parallel Nextera-based fragmentation and barcoding of individual wells. DNA from all wells is then pooled and sequenced on the Illumina HiSeq 2000 platform. Data from individual wells are demultiplexed *in silico* according to the barcode sequences. Synthetic long-reads are then assembled from the short reads using an assembly pipeline that accounts for properties of the molecular biology steps used in the library preparation (see Supplemental Materials). Because each well represents DNA from only *∼*200 molecules, even identical repeats can be resolved into synthetic reads as long as they are not so abundant in the genome as to be represented multiple times within a single well.

We applied TruSeq synthetic long-read technology to the fruit fly *D. melanogaster*, a model organism with a high quality reference genome, including extensive repeat annotation [29–31]. The version of the reference genome assembly upon which our analysis is based (Release 5.56; ftp://ftp.flybase.net/genomes/Drosophila_melanogaster/dmel_r5.56_FB2014_02/fasta/dmel-all-chromosome-r5.56.fasta.gz) contains a total of 168.7 Mbp of sequence. For simplicity, our study uses the same naming conventions as the reference genome sequence, where the sequences of chromosome arms X, 2L, 2R, 3L, 3R, and 4 contain all of the euchromatin and part of the centric heterochromatin. The sequences labelled XHet, 2LHet, 2RHet, 3LHet, 3RHet, and YHet represent scaffolds from heterochromatic regions that have been localized to chromosomes, but have not been joined to the rest of the assembly. Some of these sequences are ordered, while others are not, and separate scaffolds are separated by stretches of N’s with an arbitrary length of 100 bp. Meanwhile, the genome release also includes 10.0 Mbp of additional heterochromatic scaffolds (U) which could not be mapped to chromosomes, as well as 29.0 Mbp of additional small scaffolds that could not be joined to the rest of the assembly (Uextra). Because the Uextra sequences are generally lower quality and partially redundant with respect to the other sequences, we have excluded them from all of our analyses of assembly quality. Assembly assessment based on comparison to the Het and U sequences should also be interpreted with caution, as alignment breaks and detected mis-assemblies will partially reflect the incomplete nature of these portions of the reference sequence. Finally, we extracted the mitochondrial genome of the sequenced strain from positions 5,288,527-5,305,749 of reference sequence U using BEDTools (version 2.19.1), replacing the mitochondrial reference sequence included with Release 5.56, which represents a different strain (see http://bergmanlab.smith.man.ac.uk/?p=2033).

Approximately 50 adult individuals from the *y;cn,bw,sp* strain of *D. melanogaster* were pooled for the isolation of high molecular weight DNA, which was used to generate TruSeq synthetic long-read libraries using the aforementioned protocol (Figure S1). The strain *y;cn,bw,sp* is the same strain which was used to generate the *D. melanogaster* reference genome [25]. The fact that the strain is isogenic not only facilitates genome assembly in general, but also ensures that our analysis of TE assembly is not confounded by TE polymorphism. A total of 955,836 synthetic long-reads exceeding 1.5 Kbp (an arbitrary length cutoff) were generated with six libraries (Table S1), comprising a total of 4.20 Gbp. Synthetic long-reads averaged 4,394 bp in length, but have a local maximum near 8.5 Kbp, slightly smaller than the *∼*10 Kbp DNA fragments used as input for the protocol (Figure 1A).

**Figure 1.**
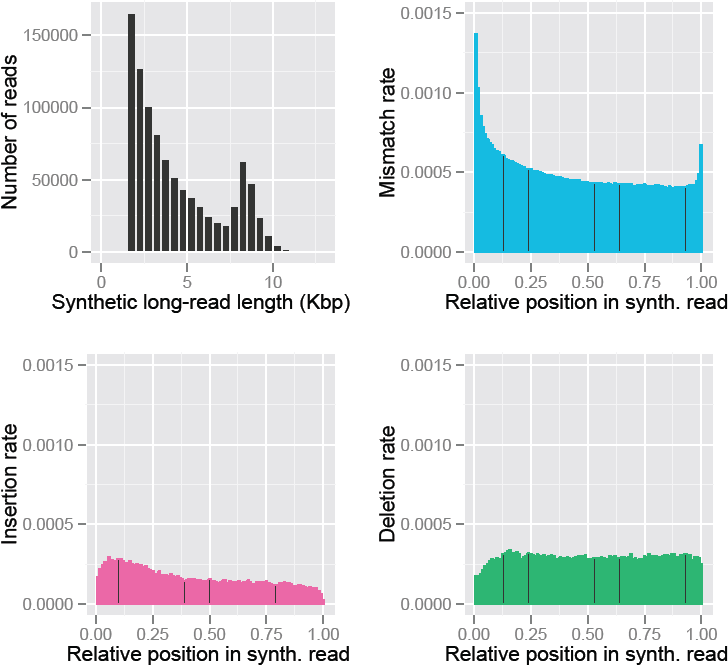
Characteristics of TruSeq synthetic long-reads. **A:** Read length distribution. **B, C, & D:** Position-dependent profiles of **B:** mismatches, **C:** insertions, and **D:** deletions compared to the reference genome. Error rates presented in these figures represent all differences with the reference genome, and can be due to errors in the reads, mapping errors, errors in the reference genome, or accurate sequencing of residual polymorphism.

#### Error rates

In order to evaluate the accuracy of TruSeq synthetic long-reads, we mapped sequences to the reference genome of *D. melanogaster*, identifying differences between the mapped synthetic reads and the reference sequence. Of 955,836 input synthetic long-reads, 99.84% (954,276 synthetic reads) were successfully mapped to the reference genome, with 90.88% (868,685 synthetic reads) mapping uniquely and 96.36% (921,090 synthetic reads) having at least one alignment with a MAPQ score *≥*20. TruSeq synthetic long-reads had very few mismatches to the reference at 0.0509% per base (0.0448% for synthetic reads with MAPQ *≥*20) as well as a very low insertion rate of 0.0166% per base (0.0144% for synthetic reads with MAPQ *≥*20) and a deletion rate of 0.0290% per base (0.0259% for synthetic reads with MAPQ *≥*20). Error rates estimated with this mapping approach are conservative, as residual heterozygosity in the sequenced line mimics errors. We therefore used the number of mismatches overlapping known SNPs to calculate a corrected error rate of 0.0286% per base (see Methods). Along with this estimate, we also estimated that the sequenced strain still retains 0.0550% residual heterozygosity relative to the time that the line was established. We note that TruSeq synthetic long-reads achieve such low error rates due to the fact that they are built as a consensuses of underlying Illumina short reads, which have an approximately ten times higher error rate. We further observed that mismatches are more frequent near the beginning of synthetic long-reads, while error profiles of insertions and deletions are relatively uniform (Figures 1B, 1C, & 1D). Minor imprecision in the trimming of adapter sequence and the error distribution along the lengths of the underlying short reads are likely responsible for this distinct error profile. Based on the observation of low error rates, no pre-processing steps were necessary in preparation for assembly, though overlap-based trimming and detection of chimeric and spurious reads are performed by default by the Celera Assembler.

#### Analysis of coverage

We quantified the average depth of coverage of the mapped synthetic long-reads for each reference chromosome arm. We observed 33.3-35.2× coverage averages of the euchromatic chromosome arms of each major autosome (2L, 2R, 3L, 3R; Figure 2). Coverage of the heterochromatic scaffolds of the major autosomes (2LHet, 2RHet, 3LHet, 3RHet) was generally lower (24.8-30.6×), and also showed greater coverage heterogeneity than the euchromatic reference sequences. This is explained by the fact that heterochromatin has high repeat content relative to euchromatin, making it more difficult to assemble into synthetic long reads. Nevertheless, the fourth chromosome had an average coverage of 34.4×, despite the enrichment of heterochromatic islands on this chromosome [32]. Depth of coverage on sex chromosomes was expected to be lower: 75% relative to the autosomes for the X and 25% relative to the autosomes for the Y, assuming equal numbers of males and females in the pool. Observed synthetic long-read depth was lower still for the X chromosome (21.2×) and extremely low for the Y chromosome (3.84×), which is entirely heterochromatic. Synthetic long-read depth for the mitochondrial genome was also relatively low (19.1×) in contrast to high mtDNA representation in short read genomic libraries, which we suspect to be a consequence of the fragmentation and size selection steps of the library preparation protocol.

**Figure 2.**
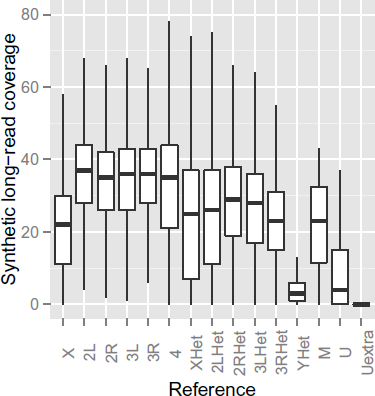
Depth of synthetic long-read coverage per chromosome arm. The suffix “Het” indicates the heterochromatic portion of the corresponding chromosome. M refers to the mitochondrial genome of the *y;cn,bw,sp* strain. U and Uextra are additional scaffolds in the reference assembly that could not be mapped to chromosomes.

### Assessment of assembly content and accuracy

#### Assembly length and genome coverage metrics

To perform *de novo* assembly, we used the Celera Assembler (version 8.1) [13], an overlap-layout-consensus assembler developed and used to reconstruct the first genome sequence of a multicellular organism, *D. melanogaster* [25], as well as one of the first diploid human genome sequences [33]. Our Celera-generated assembly contained 6,617 contigs of lengths ranging from 1,506 bp to 567.5 Kbp, with an N50 contig length of 64.1 Kbp. Note that because the TruSeq synthetic long-read data are effectively single end reads, only contig rather than scaffold metrics are reported. The total length of the assembly (i.e. the sum of all contig lengths) was 152.2 Mbp, with a GC content of 42.18% (compared to 41.74% GC content in the reference genome). Upon aligning contigs to the reference genome with NUCmer [34, 35], we observed that the ends of several contigs overlapped with long stretches (>1 Kbp) of perfect sequence identity. We therefore used the assembly program Minimus2 [36] to merge across these regions to generate supercontigs. All statistics in the following sections are based on this two-step assembly procedure combining Celera and Minimus2. The merging step resulted in the additional merging of 1,652 input contigs into 633 supercontigs, resulting in an improved assembly with a total of 5,598 contigs spanning a total of 147.4 Mbp and an N50 contig length of 69.7 Kbp (Table 1).

**Table 1.** Size and correctness metrics for de *novo* assembly. The N50 length metric measures the length of the contig for which 50% of the total assembly length is contained in contigs of that size or larger, while the L50 metric is the rank order of that contig if all contigs are ordered from longest to shortest. NG50 and LG50 are similar, but based on the expected genome size of 180 Mbp rather than the assembly length. QUAST [37] metrics are based on alignment of contigs to the euchromatic reference chromosome arms (which also contain most of the centric heterochromatin). NA50 and LA50 are analogous to N50 and L50, respectively, but in this case the lengths of aligned blocks rather than contigs are considered.

| Metric | Value |
| --- | --- |
| Number of contigs | 5598 |
| Total size of contigs | 147445959 |
| Longest contig | 567504 |
| Shortest contig | 1506 |
| Number of contigs > 10 Kbp | 2805 |
| Number of contigs > 100 Kbp | 331 |
| Mean contig size | 26339 |
| Median contig size | 10079 |
| N50 contig length | 69692 |
| L50 contig count | 554 |
| NG50 contig length | 48552 |
| LG50 contig count | 833 |
| Contig GC content | 42.26% |
| Genome fraction | 96.86% (92.24%) |
| Duplication ratio | 1.15 (1.14) |
| NA50 | 60103 (63010) |
| LA50 | 623 (618) |
| Mismatches per 100 Kbp | 7.77 (21.9) |
| Short indels ($\leq 5$ bp) per 100 Kbp | 5.10 (7.93) |
| Long indels ($> 5$ bp) per 100 Kbp | 0.46 (1.05) |
| Fully unaligned contigs | 377 (179) |
| Partially unaligned contigs | 1214 (70) |

\*Values in parentheses represent metrics calculated upon inclusion of the heterochromatic reference scaffolds (XHet, 2LHet, 2RHet, 3LHet, 3RHet, YHet, and U), which contain gaps of arbitrary size and are in some cases not oriented with respect to one another (see Release Notes at http://www.fruitfly.org/data/sequence/README.RELEASE5). Values outside of parentheses represent comparison of the assembly only to high quality reference scaffolds X, 2L, 2R, 3L, 3R, and 4.

We used the program QUAST [37] to evaluate the quality of our assembly based on alignment to the high quality reference genome. This program analyzes the NUCmer [34, 35] alignment to generate a reproducible summary report that quantifies alignment length and accuracy, as well as cataloging mis-assembly events for further investigation. Key results from the QUAST analysis are reported in Table 1, while the mis-assembly event list is included as supplemental material. The NA50 (60.1 Kbp; 63.0 Kbp upon including heterochromatic reference scaffolds) is a key metric from this report that is analogous to N50, but considers lengths of alignments to the reference genome rather than the lengths of the contigs. Contigs are effectively broken at the locations of putative mis-assembly events, including translocations and relocations. As with the synthetic long-reads, the QUAST analysis revealed that indels and mismatches in the assembly are rare, each occurring fewer than an average of 10 times per 100 Kbp (Table 1).

To gain more insight about the alignment on a per-chromosome basis, we further investigated the NUCmer alignment of the 5,598 assembled contigs to the reference genome. Upon requiring high stringency alignment (>99% sequence identity and >1 Kbp aligned), there were 3,717 alignments of our contigs to the euchromatic portions of chromosomes X, 2, 3, and 4, covering a total of 116.2 Mbp (96.6%) of the euchromatin (Table 2). For the heterochromatic sequence (XHet, 2Het, 3Het, and YHet), there were 817 alignments at this same threshold, covering 8.2 Mbp (79.9%) of the reference. QUAST also identified 179 fully unaligned contigs ranging in size from 1,951 to 26,663 bp, which we investigated further by searching the NCBI nucleotide database with BLASTN [38]. Of these contigs, 151 had top hits to bacterial species also identified in the underlying long-read data (Supplemental Materials; Table S2), 113 of which correspond to acetic acid bacteria that are known *Drosophila* symbionts. The remaining 27 contigs with no significant BLAST hit will require further investigation to determine whether they represent novel fly-derived sequences (Table S6).

**Table 2.** Alignment statistics for Celera Assembler contigs aligned to the reference genome. Alignment was performed with NUCmer [34, 35], filtering to extract only the optimal placement of each draft contig on the reference (see Supplemental Materials). Note that the number of gaps can be substantially fewer than the number of aligned contigs because alignments may partially overlap or be perfectly adjacent with respect to the reference. The number of gaps can also exceed the number of aligned contigs due to multiple partial alignments of contigs to the reference sequence.

| Reference | Aligned contigs | Alignment gaps | Length aligned (bp) | Percent aligned |
| --- | --- | --- | --- | --- |
| X | 1141 | 797 | 20720725 | 92.4% |
| 2L | 547 | 271 | 22354714 | 97.1% |
| 2R | 586 | 291 | 20645481 | 97.6% |
| 3L | 712 | 349 | 23835623 | 97.1% |
| 3R | 657 | 304 | 27453817 | 98.3% |
| 4 | 74 | 40 | 1232723 | 91.2% |
| XHet | 32 | 8 | 153247 | 75.1% |
| 2LHet | 41 | 10 | 278753 | 75.6% |
| 2RHet | 278 | 68 | 2497813 | 75.9% |
| 3LHet | 206 | 75 | 2233661 | 87.4% |
| 3RHet | 231 | 74 | 2100876 | 83.5% |
| YHet | 29 | 38 | 151545 | 43.7% |
| M | 0 | 1 | 0 | 0% |
| U | 1158 | 1198 | 4512500 | 44.9% |

#### Assessment of gene sequence assembly

In order to further assess the presence or absence as well as the accuracy of the assembly of various genomic features, we developed a simple pipeline that reads in coordinates of generic annotations and compares the reference and assembly for these sequences (see Methods). As a first step in the pipeline, we again used the filtered NUCmer [34, 35] alignment, which consists of the best placement of each draft sequence on the high quality reference genome. We then tested whether both boundaries of a given genomic feature were present within the same aligned contig. For features that met this criterion, we performed local alignment of the reference sequence to the corresponding contig using BLASTN [38], evaluating the results to calculate the proportion of the sequence aligned as well as the percent identity of the alignment. We determined that 15,684 of 17,294 (90.7%) FlyBase-annotated genes have start and stop boundaries contained in a single aligned contig within our assembly. A total of 14,558 genes (84.2%) have their entire sequence reconstructed with perfect identity to the reference sequence, while 15,306 genes have the entire length aligned with >99% sequence identity. The presence of duplicated and repetitive sequences in introns complicates gene assembly and annotation, potentially causing genes to be fragmented. For the remaining 1,610 genes whose boundaries were not contained in a single contig, we found that 1,235 were partially reconstructed as part of one or more contigs.

#### Assessment of assembly gaps

Coverage gaps and variability place an upper bound on the contiguity of genome assemblies. Regions of low coverage in synthetic long-reads may arise from biases in library preparation, sequencing, or the computational processing and assembly from underlying short read data. We therefore performed a simulation of the assembly based solely on breaks introduced by coverage gaps in the synthetic long read alignment to the reference genome (see Supplemental Materials). We required at least one overlap of at least 800 bp in order to merge across synthetic long-reads (thereby simulating a key assembly parameter) and excluded any contiguous covered region (analogous to a contig) of less than 1,000 bp. The resulting pseudo-assembly was comprised of 3,678 contigs spanning a total of 130.4 Mbp with an N50 of 80.5 Kbp. Because this expectation is based on mapping to the current reference genome, the total assembly length cannot be greater than the length of the reference sequence. Together, this simulation suggested that regions of low coverage in synthetic long-reads were primarily responsible for observed cases of assembly failure.

We next analyzed the content of the 3,524 gaps in the NUCmer alignment, which together represent failures of sequencing, library preparation, and genome assembly. We observed that 3,265 of these gaps in the whole genome assembly corresponded to previously-identified reductions in synthetic long-read coverage. Motivated by the observation that 93% of assembly gaps are explained by a deficiency in synthetic long-read coverage, we performed joint analysis of synthetic long-read and underlying short read data to help distinguish library preparation and sequencing biases from biases arising during the computational steps used to assemble the synthetic reads. We used BWA [39] to map underlying Illumina paired-end short read data to the reference genome, quantifying depth of coverage in the intervals of assembly gaps. While average short read coverage of the genome exceeded 1,500× (for MAPQ>0), mean short read coverage in the assembly gap regions was substantially lower at 263× (for MAPQ>0). However, when coverage was quantified for all mapped reads (including multi-mapped reads with MAPQ=0), average coverage was 1,153×, suggesting an abundance of genomic repeats in these intervals. We also observed a strong reduction in the GC content of the gaps (29.7%; 1,192 intervals with *<*30% GC content) compared to the overall GC content of the assembly (42.26%). This observation is therefore consistent with a known bias of PCR against high AT (but also high GC) fragments [40]. However, low GC content is also a feature of gene-poor and TE-rich regions [41], confounding this simple interpretation.

In order to gain further insight about the content of alignment gaps, we applied RepeatMasker (Smit, Hubley, & Green. RepeatMasker Open-4.0.5 1996-2010. <http://www.repeatmasker.org>) to these intervals, revealing that 35.19% of the gap sequence is comprised of TEs, 11.68% of satellites, 2.45% of simple repeats, and 0.21% of other low complexity sequence. These proportions of gap sequences composed of TEs and satellites exceed the overall genomic proportions (the fraction of the reference chromosome arms, excluding scaffolds in Uextra) of 15.07% and 1.12%, respectively, while the proportions composed of simple repeats and low complexity sequences are comparable to the overall genome proportions of 2.44% and 0.34%. Motivated by the overrepresentation of TEs in the gap intervals, we investigated which TE families were most responsible for these assembly failures. A total of 385 of the 3,524 assembly gaps overlapped the coordinates of annotated TEs, with young TE families being highly represented (Table S3). For example, LTR elements from the *roo* family were the most common, with 117 copies (of only 136 copies in the genome) overlapping gap coordinates. TEs from the *roo* family are long (canonical length of 9,092 bp) and recently diverged (mean of 0.0086 substitutions per base), and are therefore difficult to assemble (FlyTE database, http://petrov.stanford.edu/cgi-bin/Tlex_databases/flyTE_home.cgi). Conversely, elements of the high-copy number (2,235 copies) INE-1 family were underrepresented among gaps in the alignment, with only 84 copies overlapping gap coordinates. INE-1 elements tend to be short (611 bp canonical length) and represent older transposition with greater divergence among copies.

Manual curation of the alignment also revealed that assembly is particularly poor in regions of tandem arrangement of TE copies from the same family, a result that is expected because repeats will be present within individual wells during library preparation (Figure S4A). In contrast, assembly can be successful in regions with high-repeat density, provided that the TEs are sufficiently divergent or from different families (Figure S4B). Together, these observations about the assembly of particular TE families motivated formal investigation of the characteristics of individual TE copies and TE families that affect their assembly, as we describe in the following section.

#### Assessment of TE sequence assembly

Repeats can induce three common classes of mis-assembly. First, tandem repeats may be erroneously collapsed into a single copy. While the accuracy of TruSeq synthetic long-reads are advantageous in this case, such elements may still complicate assembly because they are likely to be present within a single molecule (and therefore a single well) during library preparation. Second, large repeats may fail to be assembled because reads do not span the repeat anchored in unique sequence, a situation where TruSeq synthetic long-reads are clearly beneficial. Finally, highly identical repeat copies introduce ambiguity into the assembly graph, which can result in breaks or repeat copies placed in the wrong location in the assembly. As TEs are diverse in their organization, length, copy number, GC content, and divergence, we decided to assess the accuracy of TE assembly with respect to each of these factors. We therefore compared reference TE sequences to the corresponding sequences in our assembly. Because a naive mapping approach could result in multiple reference TE copies mapping to the same location in the assembly, our approach was specifically designed to restrict the search space within the assembly based on the NUCmer global alignment (see Methods). Of the 5,434 TE copies annotated in the *D. melanogaster* reference genome, 4,565 (84.0%%) had both boundaries contained in a single contig of our assembly aligned to the reference genome, with 4,229 (77.8%) perfectly reconstructed based on length and sequence identity.

In order to test which properties of TE copies affected faithful reconstruction, we fit a generalized linear mixed model (GLMM) with a binary response variable indicating whether or not each TE copy was perfectly assembled. We included TE length as a fixed effect because we expected assembly to be less likely in cases where individual synthetic reads do not span the length of the entire TE copy. We also included GC content of the interval, including each TE copy and 1 Kbp flanking sequence on each side, as a fixed effect to capture library preparation biases as well as correlated aspects of genomic context (e.g. gene rich vs. gene poor). TE divergence estimates (FlyTE database, http://petrov.stanford.edu/cgi-bin/Tlex_databases/flyTE_home.cgi) were included as a predictor because low divergence (corresponding to high sequence identity) can cause TEs to be misplaced or mis-assembled. We also hypothesized that copy number (TE copies per family), could be important, because high copy number represents more opportunities for false joins which can break the assembly or generate chimeric contigs. Finally, we included a random effect of TE family, which accounts for various family-specific factors not represented by the fixed effects, such as sequence complexity. This grouping factor also accounts for pseudo-replication arising due to multiple copies of TEs within families [42]. We found that length (*b* = −1.633, *Z* = −20.766, P < 2 × 10^−16^), divergence (*b* = 0.692, *Z* = 7.501, *P* = 6.35 × 10^−14^), and GC content (*b* = 0.186, *Z* = 3.171, *P* = 0.00152) were significant predictors of accurate TE assembly (Figure 3; Table S5). Longer and less divergent TE copies, as well as those in regions of low GC content, resulted in a lower probability of accurate assembly (Figure 3). We found that overall copy number was not a significant predictor of accurate assembly (*b* = 0.095, *Z* = 0.162, *P* = 0.871). However, upon restricting the test to consider only high identity copies (*<*0.01 substitutions per base compared to the canonical sequence), we observed an expected reduction in the probability of accurate assembly with increasing copy number (*b* = −0.529, *Z* = −2.936, *P* = 0.00333). Plotting initial results also suggested a possible interaction between divergence and the number of high identity copies. Our model therefore additionally includes this significant interaction term, which demonstrates that low divergence of an individual TE copy is more problematic in the presence of many high identity copies from the same family (Figure 3).

**Figure 3.**
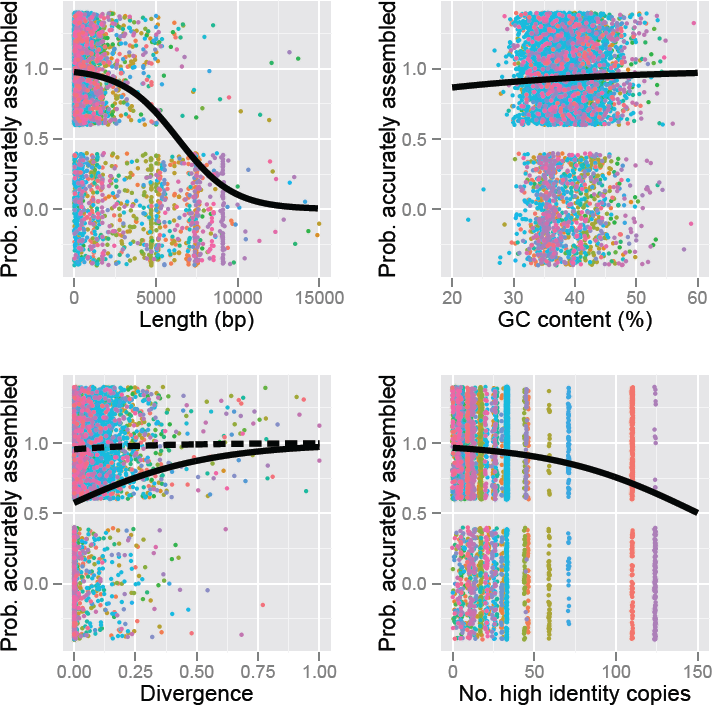
Results of generalized linear mixed model describing probability of accurate TE assembly. Predictor variables include: TE length (*b* = −1.633, *Z* = −20.766, *P* < 2 × 10^−16^), GC content (*b* = 0.186, *Z* = 3.171, *P* = 0.00152), divergence (*b* = 0.692, *Z* = 7.501, *P* = 6.35 × 10^−14^), and number of high identity (*<* 0.01 substitutions per base compared to the canonical sequence) copies within family (*b* = −0.529, *Z* = −2.936, *P* = 0.00333). Black lines represent predicted values from the GLMM fit to the binary data (colored points). The upper sets of points represent TEs which were perfectly assembled, while the lower set of points represent TEs which are absent from the assembly or were mis-assembled with respect to the reference. The exact positions of the colored points along the Y-axis should therefore be disregarded. Colors indicate different TE families (122 total). To visualize the interaction between divergence and the number of high identity copies (*b* = 0.382, *Z* = 3.921, *P* = 8.81 × 10^−5^), we plotted predicted values for both families with low numbers of high identity copies (dashed line) as well as families with high numbers of high identity copies (solid line).

In spite of the limitations revealed by our analysis, we observed several remarkable cases where accurate assembly was achieved, distinguishing the sequences of TEs from a single family with few substitutions among the set. For example, elements in the *Tc1* family have an average of 0.039 substitutions per base with respect to the 947 bp canonical sequence, yet 25 of 26 annotated copies were assembled with 100% accuracy (Table S4). The assembled elements from this family range from 131 bp to 1,662 bp, with a median length of 1,023 bp.

### Impact of the coverage on assembly results

The relationship between coverage and assembly quality is complex, as we expect a plateau in assembly quality at the point where the assembly is no longer limited by data quantity. To evaluate the impact of depth of synthetic long-read coverage on the quality of the resulting assembly, we randomly down-sampled the full *∼*34× dataset to 20×, 10×, 5×, and 2.5×. We then performed separate *de novo* assemblies for each of these down-sampled datasets, evaluating and comparing assemblies using the same size and correctness metrics previously reported for the full-coverage assembly. We observed an expected nonlinear pattern for several important assembly metrics, which begin to plateau as data quantity increases. NG50 contig length (analogous to N50, but normalized to the genome size of 180 Mb to facilitate comparison among assemblies) increases rapidly with coverage up to approximately 10×, increasing only marginally at higher synthetic long-read coverage (Figure 4A). We do not expect the monotonic increase to continue indefinitely, as very high coverage can overwhelm OLC assemblers such as Celera (see documentation, which advises against high coverage such as 80×). Gene content of the assembly also increases only marginally as synthetic long-read coverage increases above approximately 10×, but TE content does not saturate as rapidly (Figure 4B). Our results likewise suggest that even very low synthetic long-read coverage assemblies (5×) can accurately recover approximately half of all genes and TEs.

**Figure 4.**
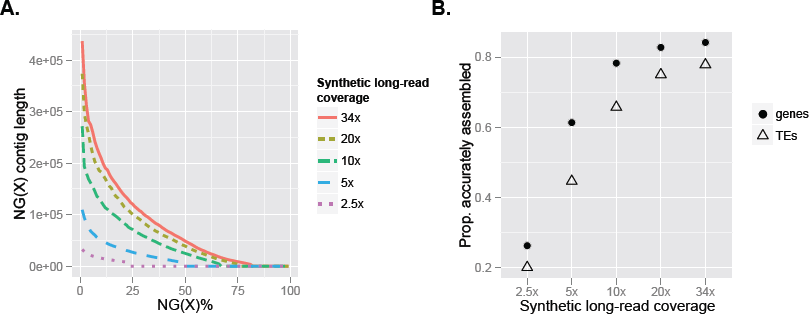
Assembly metrics as a function of depth of coverage of TruSeq synthetic long-reads. **A:** NG(X) contig length for full and down-sampled coverage data sets. This metric represents the size of the contig for which X% of the genome length (180 Mbp) lies in contigs of that size or longer. **B:** The proportion of genes and transposable elements accurately assembled (100% length and sequence identity) for full and down-sampled coverage data sets.

## Discussion

Rapid technological advances and plummeting costs of DNA sequencing technologies have allowed biologists to explore the genomes of species across the tree of life. However, translating the massive amounts of sequence data into a high quality reference genome ripe for biological insight is a substantial technical hurdle. Many assemblers use coverage-based heuristics to classify problematic repeats and either break the assembly at ambiguous repeat regions or place consensus repeat sequences in the assembly. This approach balances the tradeoff between assembly contiguity and the rate of mis-assembly, but the resulting biased representation of certain classes of repeats limits understanding of repeat evolution. Understanding the dynamics of repeats such as TEs is fundamental to the study genome evolution, as repeats affect genome size structure as well as genome function [7, 9, 43–45]. Several tools (e.g. T-lex2 [46], RetroSeq [47], Tea [48], ngs_te_mapper [49], RelocaTE [50], RetroSeq [47], PoPoolation TE [51], TE-locate [52]) are currently available for discovery and annotation of TE sequences in high-throughput sequencing data. However, because these tools depend on the quality of the assembly to which they are applied, annotation is generally limited to TE families containing predominantly short and divergent TE copies, biasing our current view of TE organization. Accurate assembly and annotation of TEs and other repeats will dramatically enrich our understanding of the complex interactions between TEs and host genomes as well as genome evolution in general.

One of the simplest ways to accurately resolve repeat sequences is to acquire reads longer than the lengths of the repeats themselves. Here, we evaluated a novel library preparation approach that allows the generation of highly accurate synthetic reads up to 18.5 Kbp in length. We tested the utility of this approach for assembling and placing highly repetitive, complex TEs with high accuracy. As a first step in our analysis, we analyzed the content of the synthetic long-read data, evaluating synthetic long-read accuracy as well as coverage of the *D. melanogaster* reference genome. We found that the synthetic long-reads were highly accurate, with error rates comparable to consensus sequences produced using third generation long-read sequencing technologies. We also observed relatively uniform coverage across both the euchromatic and heterochromatic portions of the autosomes, with an expected reduced coverage of the heterochromatin. This observation is explained by the fact that heterochromatin is generally more repetitive and therefore more difficult to assemble into long reads from underlying short read data.

Despite general uniformity in synthetic long-read coverage, we identified important biases resulting in coverage gaps and reductions in synthetic long-read coverage in repeat-dense regions with relatively low average GC content. While GC biases of PCR are well documented, GC content is also correlated with repeat density, thereby confounding this interpretation [41]. Other biases introduced in the molecular biology, sequencing, and/or computational steps of the data preparation (e.g. the fragility of certain DNA sequences during the random shearing step) are also possible, but cannot be disentangled using this data set and will require further investigation. Enhancements to the protocol enabled by a better understanding of these biases could substantially improve the utility of the technology, as reductions in synthetic long-read coverage explained the vast majority of gaps in the genome assembly.

Our assembly achieved an N50 contig length of 69.7 Kbp, covering 96.9% of the euchromatic scaffolds (and centric heterochromatin) of the reference genome, and containing 84.2% of annotated genes with perfect sequence identity. Using standard assembly size (number of contigs, contig length, etc.) and correctness metrics based on alignment to the reference genome, we demonstrated that our assembly is comparable to other *de novo* assemblies of large and complex genomes [53, 54]. Nevertheless, we expect that future methodological advances will unlock the full utility of TruSeq synthetic long-read technology. We used a simple pipeline of existing tools to investigate the advantages and limitations of TruSeq synthetic long-reads, but new algorithms and assembly software will be tailored specifically for this platform in the near future (J. Simpson, pers. comm.).

An important caveat in the interpretation of our results is the fact that the assembly was performed on a highly inbred strain of *D. melanogaster*. This was beneficial to our study because it allowed us to attribute TE sequence differences to divergence among TE copies. For an outbred species, distinguishing between divergence among TE copies and polymorphism within TE copies complicates this analysis. For the same reasons, polymorphism in general is a key feature limiting non-haploid genome assemblies, as algorithms must strike a balance between merging polymorphic haplotypes and splitting slightly diverged repeat copies to produce a haploid representation of the genome sequence. Forthcoming assemblies of other *Drosophila* from the modENCODE project (using 454 reads) demonstrate the importance of polymorphism, achieving N50 contig lengths up to 436 Kbp for inbred species, but only 19 Kbp for the species that could not be inbred [55]. The first application of the synthetic long-read technology presented here was to assemble the genome of the colonial tunicate *Botryllus schlosseri*, but assessment of assembly quality was difficult as no high quality reference genome exists for comparison. Likewise, recent work demonstrated the utility of the same technology for assigning polymorphisms to individual haplotypes [56], but this problem is somewhat distinct from the *de novo* resolution of polymorphism in the absence of a reference genome. Future work will be required to systematically evaluate the ability of synthetic long-read data to help resolve polymorphism in outbred species.

Our study demonstrates that TruSeq synthetic long-reads enable accurate assembly of complex, highly repetitive TE sequences. Previous approaches to *de novo* assembly generally fail to assemble and place long, abundant, and identical TE copies with respect to the rest of the assembly. For example, the majority of TE-containing contigs in the improved draft assembly of *Drosophila simulans* (which combined Illumina short read and Sanger data) were smaller than 500 bp [16]. Likewise, short read assemblies from the *Drosophila* 12 Genomes Consortium [15] estimated TE copy number, but did not even attempt to place TE sequences with respect to the rest of the assemblies. Our assembly contains 77.8% of annotated TEs perfectly identical in sequence to the current reference genome. Despite the high quality of the current reference, errors undoubtedly exist in the current TE annotations, and it is likely that there is some divergence between the sequenced strain and the reference strain from which it was derived, making our estimate of the quality of TE assembly conservative. Likewise, we used a generalized linear modeling approach to demonstrate that TE length is the main feature limiting the assembly of individual TE copies, a limitation that could be partially overcome by future improvements to the library preparation technology to achieve even longer synthetic reads. This analysis also revealed a significant interaction between divergence and the number of high identity copies within TE families. Low divergence among copies is problematic for families with a large number of high identity copies, but is less important for families with few high identity copies. Further dilution during library preparation may therefore enhance assembly of dispersed TE families. By performing this assessment in *D. melanogaster*, a species with particularly active, abundant, and identical TEs, our results suggest that synthetic long-read technology can empower studies of TE dynamics for many non-model species.

Alongside this synthetic long-read technology, several third-generation sequencing platforms have been developed to sequence long molecules directly. One such technology, Oxford Nanopore (Oxford, UK) sequencing [21], possesses several advantages over existing platforms, including the generation of reads exceeding 5 Kbp at a speed of 1 bp per nanosecond. Pacific Biosciences’ (Menlo Park, CA, USA) single-molecule real-time (SMRT) sequencing platform likewise uses direct observation of enzymatic reactions to produce base calls in real time with reads averaging *∼*8.5 Kbp in length (for P5-C3 chemistry), and fast sample preparation and sequencing (1-2 days each) [57, http://investor.pacificbiosciences.com/releasedetail.cfm?ReleaseID=794692]. Perhaps most importantly, neither Nanopore nor PacBio sequencing requires PCR amplification, thereby reducing biases and errors that place an upper limit on the sequencing quality of most other platforms. By directly sequencing long molecules, these third-generation technologies will likely outperform TruSeq synthetic long-reads in certain capacities, such as assembly contiguity enabled by homogeneous genome coverage. Indeed, preliminary results from the assembly of a different *y;cn,bw,sp* substrain of *D. melanogaster* using corrected PacBio data achieved an N50 contig length of 15.3 Mbp and closed two of the remaining gaps in the euchromatin of the Release 5 reference sequence (Landolin et al., 2014 [http://dx.doi.org/10.6084/m9.figshare.976097]). While not yet systematically assessed, it is likely that PacBio long reads will also help resolve high identity repeats, though current raw error rates may be limiting.

Most current approaches to *de novo* assembly fare poorly on long, abundant, and recently diverged repetitive elements, including some families of TEs. The resulting assemblies offer a biased perspective of evolution of complex genomes. In addition to accurately recovering 96.9% of the euchromatic portion of the high quality reference genome, our assembly using TruSeq synthetic long-reads accurately placed and perfectly reconstructed the sequence of 84.2% of genes and 77.8% of TEs. Improvements to *de novo* assembly, facilitated by TruSeq synthetic long-reads and other long read technologies, will empower comparative analyses that will enlighten the understanding of the dynamics of repeat elements and genome evolution in general.

## Materials and Methods

### Reference genome and annotations

The latest release of the *D. melanogaster* genome sequence at the time of the preparation of this manuscript (Release 5.56) and corresponding TE annotations were downloaded from FlyBase (http://www.fruitfly.org/). All TE features come from data stored in the FlyTE database (http://petrov.stanford.edu/cgi-bin/Tlex_databases/flyTE_home.cgi), and were detected using the program BLASTER [30, 31].

### Library preparation

High molecular weight DNA was separately isolated from pooled samples of the *y;cn,bw,sp* strain of *D. melanogaster* using a standard ethanol precipitation-based protocol. Approximately 50 adult individuals, both males and females, were pooled for the extraction to achieve sufficient gDNA quantity for preparation of multiple TruSeq synthetic long-read libraries.

Six libraries were prepared by Illumina’s FastTrack Service using the TruSeq synthetic long-read technology, previously known as Moleculo. To produce each library, extracted gDNA is sheared into approximately 10 Kbp fragments, ligated to amplification adapters, and then diluted to the point that each well on a 384-well plate contains approximately 200 molecules, representing approximately 1.5% of the entire genome. These pools of DNA are then amplified by long range PCR. Barcoded libraries are prepared within each well using Nextera-based fragmentation and PCR-mediated barcode and sequencing adapter addition. The libraries undergo additional PCR amplification if necessary, followed by paired-end sequencing on the Illumina HiSeq 2000 platform.

### Assembly of synthetic long-reads from short read data

Based on the unique barcodes, assembly is performed among molecules originating from a single well, which means that the likelihood of individual assemblies containing multiple members of gene families (that are difficult to distinguish from one another and from polymorphism within individual genes) is greatly reduced. The assembly process, which is described in detail in the Supplemental Materials, consists of several modules. First, raw short reads are pre-processed to remove low-quality sequence ends. Digital normalization [58] is then performed to reduce coverage biases introduced by PCR, such that the corrected short read coverage of the highest covered fragments is *∼*40×. The next step uses overlap-based error correction to generate higher quality consensus sequences for each short read. The main assembly steps implement the String Graph Assembler (SGA) [59] which generates contigs using an overlap approach, then scaffolds contigs from the same fragment using paired-end information. Gap filling is then conducted to fill in scaffold gaps. The original paired-end reads are then mapped back to the assembled synthetic long-reads and contigs are either corrected or broken based on inconsistencies in the alignment.

### Assessment of synthetic long-read quality

To estimate the degree of contamination of the *D. melanogaster* libraries prepared by Illumina, we used BLASTN (version 2.2.28+) [38] to compare the synthetic long-reads against reference sequences from the NCBI nucleotide database (http://www.ncbi.nlm.nih.gov/nuccore), selecting the target sequences with the lowest e-value for each query sequence.

The TruSeq synthetic long-reads were then mapped to the *D. melanogaster* reference genome as single-end reads using BWA-MEM [39]. Depth of coverage was estimated by applying the GATK DepthOfCoverage tool to the resulting alignment. To estimate error rates, we then parsed the BAM file to calculate position-dependent mismatch, insertion, and deletion profiles. Because a portion of this effect would result from accurate sequencing of genomes harboring residual heterozygosity, we used data from the *Drosophila* Genetic Reference Panel (DGRP) [60] to estimate both the rate of residual heterozygosity as well as a corrected error rate of the TruSeq synthetic long-reads. We applied the jvarkit utility (<https://github.com/lindenb/jvarkit/wiki/SAM2Tsv>) to identify positions in the reference genome where mismatches occurred. We then used the relationship that the total number sites with mismatches to the euchromatic reference chromosome arms (*M*) = 1,105,831 = *Lm* + *pLθ*, where *L* is the 120,381,546 bp length of the reference sequence to which we aligned, *m* is the per base error rate, *p* is the proportion of heterozygous sites still segregating in the inbred line, and *θ* is the average proportion of pairwise differences between *D. melanogaster* genome sequences, estimated as 0.141 from DGRP. Meanwhile, the number of mismatches that overlap with SNP sites in DGRP (*M_SNP_*) = 53, 515 = *Lmθ_D_*+*pLθ*, where *θ_D_* is the proportion of sites that are known SNPs within DGRP (0.0404). Note that this formulation makes the simplifying assumption that all segregating SNPs would have been previously observed in DGRP, which makes the correction conservative. Solving for the unknown variables:

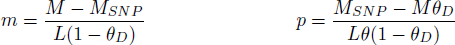

To convert *m* to the TruSeq synthetic long-read error rate, we simply divide by the average depth of coverage of the euchromatic sequence (31.81×), estimating a corrected error rate of 0.0286% per base. This estimate is still conservative in that it does not account for mismatches observed multiple times at a single site, which should overwhelmingly represent residual polymorphism.

### Genome assembly

Most recent approaches to *de novo* genome assembly are based on the de Bruijn graph paradigm, which offers a substantial computational advantage over overlap-layout-consensus (OLC) approaches when applied to large datasets. Nevertheless, for datasets with moderate sequencing depth (such as TruSeq synthetic long-read libraries), OLC approaches can be computationally tractable and tend to be less affected by both repeats and sequencing errors than de Bruijn graph-based algorithms. Likewise, many modern Bruijn graph-based assemblers simply do not permit reads exceeding arbitrary length cutoffs. We therefore elected to use the Celera Assembler [13], an OLC assembler developed and used to generate the first genome sequence of a multicellular organism, *Drosophila melanogaster* [25], as well as one of the first diploid human genome sequences [33].

As TruSeq synthetic long-reads share some characteristics with consensus-corrected PacBio reads, we applied Celera Assembler parameters recommended for these PacBio data to take advantage of the read length and low error rate [23, 61]. In particular, the approach uses a different unitigger algorithm, decreases the unitig error rates (which is made possible by low synthetic long-read error rates and low rates of polymorphism) and increased the k-mer size to increase overlap specificity. Upon observing partially overlapping contigs among the output of the Celera Assembler, we decided to use the program Minimus2 [36] to merge these contigs into supercontigs, reducing redundancy and improving assembly contiguity. Parameters used for both assembly programs are further described in the Supplemental Materials.

For the down-sampled assemblies with lower coverage, we based the expected coverage on the average euchromatic autosomal depth of coverage of 34× for the full synthetic long-read dataset. We randomly sampled reads from a concatenated FASTQ of all six libraries until the total length of the resulting dataset was equal to the desired coverage.

### Assessment of assembly quality

We aligned the contigs produced by the Celera Assembler to the reference genome sequence using the NUCmer pipeline (version 3.23) [34, 35]. From this alignment, we used the delta-filter tool to extract the best mapping of each query draft contig onto the high quality reference sequence (see Supplemental Materials). We then used to coordinates of these alignments to both measure overall assembly quality and investigate assembly of particular genomic features, including genes and TEs. Using this alignment, we identified the locations of reference-annotated gene and TE sequences in our assembly and used local alignment with BLASTN [38] to determine sequence identity and length ratio (assembled length/reference length) for each sequence. To calculate correctness metrics, we used the tool QUAST (version 2.3) which again uses the NUCmer alignment to the reference genome to calculate the prevalence of mismatches, indels, and other mis-assembly events.

The GLMM used to test the characteristics of TEs that affected accurate assembly were built using the *lme4* package [62] within the R statistical computing environment [63]. TE features (predictor variables) were available for all but the *Y* family of TEs, which was recently annotated (Release 5.56). The response variable was represented by a binary indicator denoting whether or not the entire length of the TE was accurately assembled. This model assumed a binomial error distribution with a logit link function. TE copy length, GC content (including 1 Kbp flanking regions on each side), divergence (number of substitutions per base compared to the canonical sequence of the TE family), number of high identity (*<*0.01 substitutions per base compared to the canonical sequence) copies per family, and the interaction between high identity copies and divergence were included as fixed effects, while TE family was included as a random effect. All predictor variables were standardized to zero mean and unit variance prior to fitting, in order to compare the magnitude of the effects.

All figures with the exception of those in the supplement were generated using the *ggplot2* package [64].

### Data access

Sequence data can be found under the NCBI BioProject: PRJNA235897, BioSample: SAMN02588592. Experiment SRX447481 references the synthetic long-read data, while experiment SRX503698 references the underlying short read data. The main genome assembly is available from FigShare at http://dx.doi.org/10.6084/m9.figshare.985645 and the QUAST contig report is available at http://dx.doi.org/10.6084/m9.figshare.985916. Scripts written to assess presence or absence of genomic features in the *de novo* assembly can be found in a GitHub repository at https://github.com/rmccoy7541/ assess-assembly while other analysis scripts, including those to reproduce down-sampled assemblies, can be found in a separate GitHub repository at https://github.com/rmccoy7541/dmel-longread-assembly. The parameter choices for various software packages are described in the Supplemental Materials.

## Acknowledgments

Thank you to Alan Bergland for performing the DNA extractions and to Anthony Long for providing the strain. Thanks also to Julie Collens and Courtney McCormick for preparing and delivering the synthetic long-read and underlying short read libraries. And thank you to Jared Simpson for advice regarding the genome assembly, as well as the engineers and administrators of the Proclus and SCG computing clusters.

